# Effect of HSV-1 ICP0 E3 ubiquitin ligase on CIN85 endosomes

**DOI:** 10.64898/2026.09.21.753399

**Authors:** Sarah Lasnier, Maria Kalamvoki

## Abstract

Herpes simplex virus-1 (HSV-1) has infected more than 67% of the world population causing diseases that range in severity from benign cold sores to encephalitis. To infect and persist in the host, HSV-1 has evolved sophisticated strategies to counteract antiviral responses. The viral immediate early protein ICP0 (Infected Cells Protein 0) plays a fundamental role in this process. ICP0 is an E3 ubiquitin ligase that is required for successful onset of lytic infection and productive reactivation of viral genomes from latency. Most ICP0 studies have been focused on its nuclear functions, but late gene expression drives accumulation of ICP0 to the cytoplasm where its functions are unknown. We recently demonstrated that cytoplasmic ICP0 localizes on the surface of endosomes through its interaction with CIN85 (Cbl-interacting protein of 85 kDa), a scaffolding protein for factors involved in endocytosis. This interaction is required for extrusion of host antiviral factors and immunoevasion. Here, we report that the E3 ubiquitin ligase activity of ICP0 is also required for these effects, as disruption of ICP0 RING finger domain, which disrupts its catalytic activity, resulted in an increase in the number of CIN85 endosomes that cannot be exocytosed. These endosomes colocalize with autophagosome components, but they do not enter the lysosomal pathway for degradation. Mutant viruses that cannot exocytose CIN85 endosomes produce EVs that are more potent at activating antiviral responses. Overall, this study identifies the HSV-1 ICP0 E3 ubiquitin ligase as a key determinant of the fate of CIN85 endosomes, which are important for immunoevasion.

**Importance:** Herpes simplex virus (HSV-1) infection can significantly impact not only infected cells, but also neighboring and distant cells. In infected cells, HSV-1 evades antiviral responses and subverts host mechanisms to support virus replication and progeny virus production. Neighboring and distant cells are impacted both directly, by progeny viruses, and indirectly, by particles produced by infected cells. These particles are largely comprised of EVs. In infected cells, HSV-1 alters the cargo and biogenesis of EVs, influencing the microenvironment, and driving pathogenesis. Here, we report that HSV-1 controls the properties of CIN85 endosomes through its E3 ubiquitin ligase ICP0 and diverts them for exocytosis. In this way, HSV-1 extrudes host antiviral factors to evade host responses. EVs released by HSV-1-infected cells activate type-I interferon (IFN) and pro-inflammatory responses, potentially driving pathogenesis.

## Introduction

Herpes simplex virus 1 (HSV-1) has infected more than 67% of the population worldwide. HSV-1 primarily infects mucosal epithelial cells and establishes lifelong infections in sensory neurons. Periodic reactivation of the virus causes only benign vesicular lesions, but frequent reactivation, particularly in elderly and immunocompromised individuals, can cause more severe diseases (1).

HSV-1 has evolved a plethora of mechanisms to evade host antiviral responses and to subvert host machinery to support the infection. The Infected Cell Protein 0 (ICP0) plays a fundamental role in these processes. ICP0 is encoded by an immediate-early gene of the virus and is known as a promiscuous transactivator as it enables expression both of host and viral genes (2–7). To achieve this, ICP0 disperses chromatin repressor complexes from the viral genome and recruits chromatin remodeling enzymes to enable viral gene expression (8–11). ICP0 also functions as an E3 ubiquitin ligase; this activity is conserved among alphaherpesvirus ICP0 orthologues (12–17). Major ICP0 targets include components of the nuclear domain 10 (ND10) bodies, where the viral genome is transcriptionally repressed, histone chaperons, DNA damage response factors, and innate immunity components (18–30). Because of these roles, ICP0 is required *in vivo* for lytic infection and productive reactivation of viral genomes from latency (7, 14, 17, 31–38).

ICP0 displays distinct localization patterns in different stages of HSV-1 infection that facilitate its different functions. It initially localizes in the nucleus and appears punctated as it accumulates at the ND10 bodies, where it degrades key structural components and exerts its transactivation function. After the replication compartments have been formed, ICP0 displays a diffuse nuclear localization pattern. Finally, late gene expression triggers accumulation of ICP0 into the cytoplasm where it displays both diffuse and punctate pattern, but its further functions are unknown (39–42). Earlier studies discovered that ICP0 in the cytoplasm interacts with CIN85 (Cbl-interacting protein of 85 kDa), an endocytosis adaptor protein, to accelerate the removal of EGFR (epidermal growth factor receptor) from the surface of HSV-1-infected cells and suppress its signaling (40). Our lab then demonstrated that ICP0 and CIN85 remove Nectin-1 (a virus entry receptor) from the surface of the infected cells, to ensure successful virus spread (39, 40). Combined, these findings indicate a novel function for ICP0 in endocytosis, though the precise mechanism is unknown. We recently demonstrated that ICP0 interacts and colocalizes with CIN85 on the surface of vesicular structures in the cytoplasm of infected cells (43). CIN85 is comprised of three N-terminal SH3 domains, followed by a central proline-rich region and a C-terminal coiled-coil domain. CIN85 docks on vesicles via its coiled-coil domain and positively charged amino acids in its C-terminus (44). Via its SH3 domains, CIN85 binds a consensus Px(P/A)xxR motif in proteins involved in endocytosis and protein sorting (39, 40, 45–48). ICP0 has two consecutive CIN85-binding motifs between amino acids 244-277 that are necessary for its recruitment to CIN85 endosomes (40, 43). In infections with ICP0 Δ244-277 virus cytoplasmic ICP0 remains diffuse as it does not interact with CIN85 (43). The interaction of ICP0 with CIN85 is critical for exocytosis of host factors and the ability of the virus to evade host antiviral responses (43).

The interaction of ICP0 with CIN85 is essential for recruitment of ICP0 to CIN85 endosomes, however, the function of ICP0 on endosomes remains unknown. ICP0 has an E3 Ub ligase activity that is attributed to its N-terminal zinc-binding RING-finger domain (RF), located between the amino acids 116-156 of the protein (49). ICP0 has been implicated in ubiquitination and degradation of a variety of host substrates, most of which have nuclear functions (27, 30, 50). Whether ICP0 also targets substrates in the cytoplasm remains understudied. ICP0 has been implicated so far in K48-linked polyubiquitination of substrates that are targeted for proteasomal degradation, but other types of ubiquitination and different fates of substrate proteins cannot be excluded (51–55).

Here, we sought to determine whether the E3 ubiquitin ligase activity of ICP0 impacts the CIN85 endosomes and their trafficking. We observed a numerical increase in the ICP0/CIN85 vesicles after infections with a RF-ICP0 mutant virus, with impaired E3 ubiquitin ligase activity, compared to infections with the wild-type virus (WT). This phenotype was conserved across multiple cell types and primary cells. ICP0 was not observed on vesicles when CIN85 was not expressed. This is consistent with our previous findings showing that the interaction of ICP0 with CIN85 is essential for localization of ICP0 to CIN85 endosomes (43). The ICP0/CIN85 endosomes observed in RF-ICP0 infected cells are not autophagosomes and they do not associate with lysosomes, even though they colocalize with the autophagosome markers LC3 (microtubule-associated protein 1A/B light chain 3) and ATG5 (autophagy-related protein 5), and p62/SQSTM1 (sequestosome 1). Also, they are distinct from the CD63 endosomes, whose exocytosis is stimulated in HSV-1-infected cells (56–58). WT virus infection promotes exocytosis of the CIN85 endosomes, but this is not the case in RF-ICP0-infected cells or after infection with the ICP0Δ244-277 virus, where ICP0 does not interact with CIN85. Thus, our findings indicate that the binding of ICP0 to CIN85, as well as ICP0’s E3 ubiquitin ligase activity, are both necessary to promote exocytosis of CIN85 endosomes. Extracellular vesicles (EVs) released by RF-ICP0 virus infected cells mount stronger antiviral responses compared to EVs released by WT-virus infected cells, indicating that the ICP0 E3 ubiquitin ligase is used by the virus to evade the host both inside the cells as well as the surrounding microenvironment. Overall, we have provided evidence that the ICP0 E3 ubiquitin ligase activity regulates CIN85 endosomes.

## Results

### Infection with an ICP0 E3 ubiquitin ligase mutant virus (RF-ICP0) displays aberrant ICP0/CIN85 vesicles in multiple cell types

We have previously shown ICP0 and CIN85 to colocalize in vesicular structures in the cytoplasm of HSV-1 infected cells, and that recruitment of ICP0 to these vesicles depends on the two CIN85-binding motifs of ICP0 located between amino acids 244-277 (Px(P/A)xxR) (43). The E3 ubiquitin ligase of ICP0 is known to ubiquitinate host factors during WT infection for degradation. To determine whether the ICP0 E3 ubiquitin ligase impacts CIN85 endosomes, we used RF-ICP0 virus with inactive E3 ubiquitin ligase due to two point mutations, C116A and C156A, in the RING finger domain of ICP0 (59). Using this virus we performed infections (10 PFU/cell) in human adult primary epidermal keratinocytes (HEKa) and observed an increase in the ICP0 vesicles per cell compared to WT virus-infected cells by 15 h post-infection using an immunofluorescence approach (Figure 1A). We quantified these vesicles and determined that RF-ICP0 infected cells have six times as many vesicles as WT virus. As a control, we infected these cells with the ICP0Δ244-277 virus and observed that ICP0 remained diffuse in the cytoplasm (Figure 1A). Similar results we obtained in WT virus versus RF-ICP0 infections in human immortalized keratinocytes (HaCaT) (Figure 1B), hTERT-immortalized human embryonic lung fibroblasts (hTERT-HEL) (Figure 2A), and the spontaneously immortalized Adult Retinal Pigment Epithelial cell line-19 (ARPE-19) (data not shown). Finally, to determine whether these ICP0 vesicles also contain CIN85 we expressed exogenous CIN85 in HEp-2 cells followed by infection with either the WT virus or the RF-ICP0 mutant (10 PFU/cell). We observed that all ICP0 vesicles were positive for CIN85 (Figure 2B). We also observed that ICP0/CIN85 vesicles coalesce into a few larger structures in WT virus infected cells, but numerous smaller vesicles were observed in RF-ICP0 infection (Figure 2B). Overall, we discovered that disruption of the ICP0 E3 ubiquitin ligase activity increases the number of ICP0/CIN85 endosomes in infected cells.

**Figure 1:**
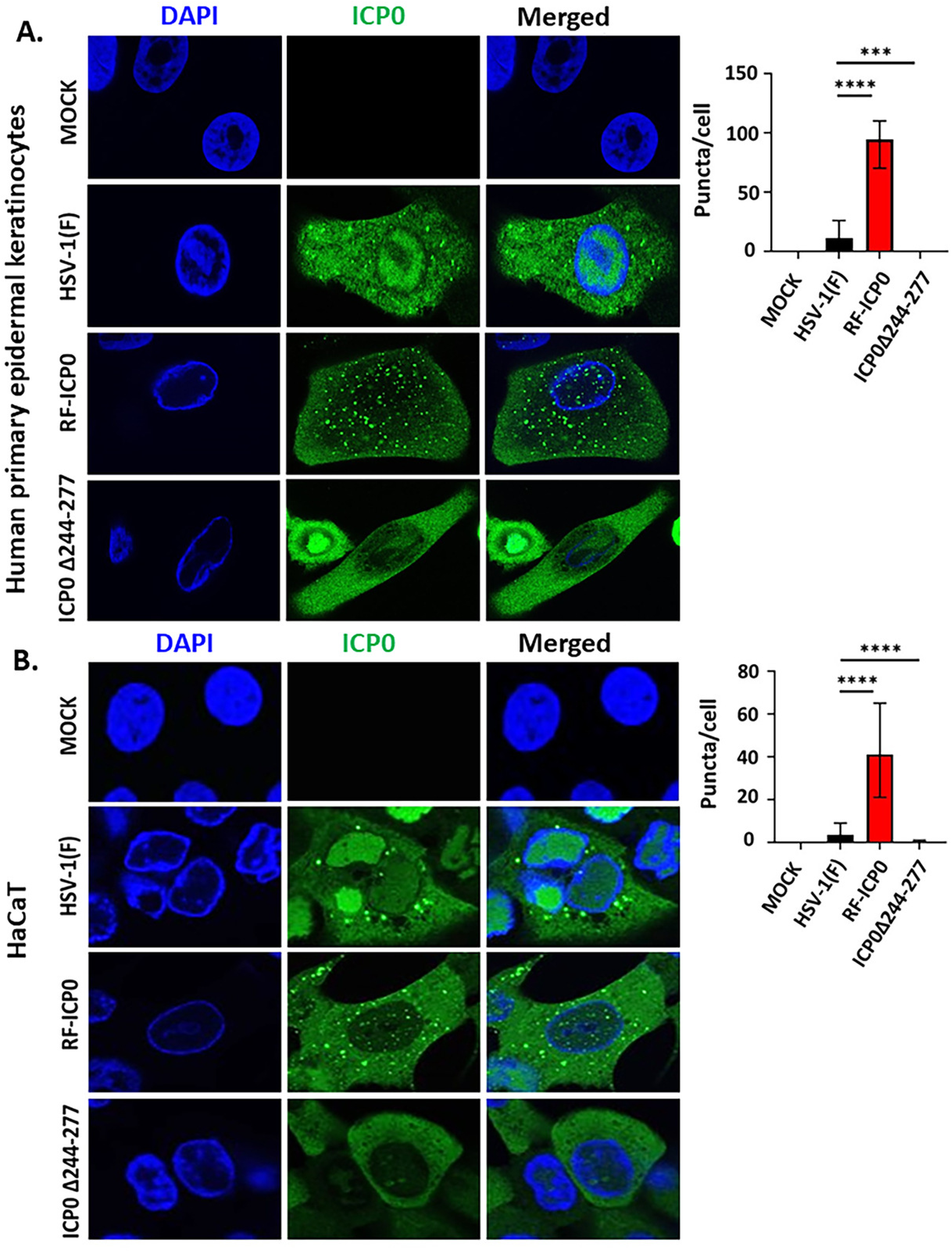
Disruption of HSV-1 ICP0 E3 ubiquitin ligase activity increases CIN85 endosomes. **(A)** Human adult primary keratinocytes (HEKa) and HaCAT **(B)** were infected with WT, RF-ICP0, and ICP0Δ244-277 virus (10 PFU/cell). Cells were fixed at 15 h post-infection and stained with ICP0 antibody. Cell nuclei were stained with DAPI. Images were captured using a Leica TCS SP8 confocal microscope. Puncta numbers per cell were quantified using Image J from approximately 20 cells from 3 independent experiments. Graphs were made using GraphPad Prism 10.

**Figure 2:**
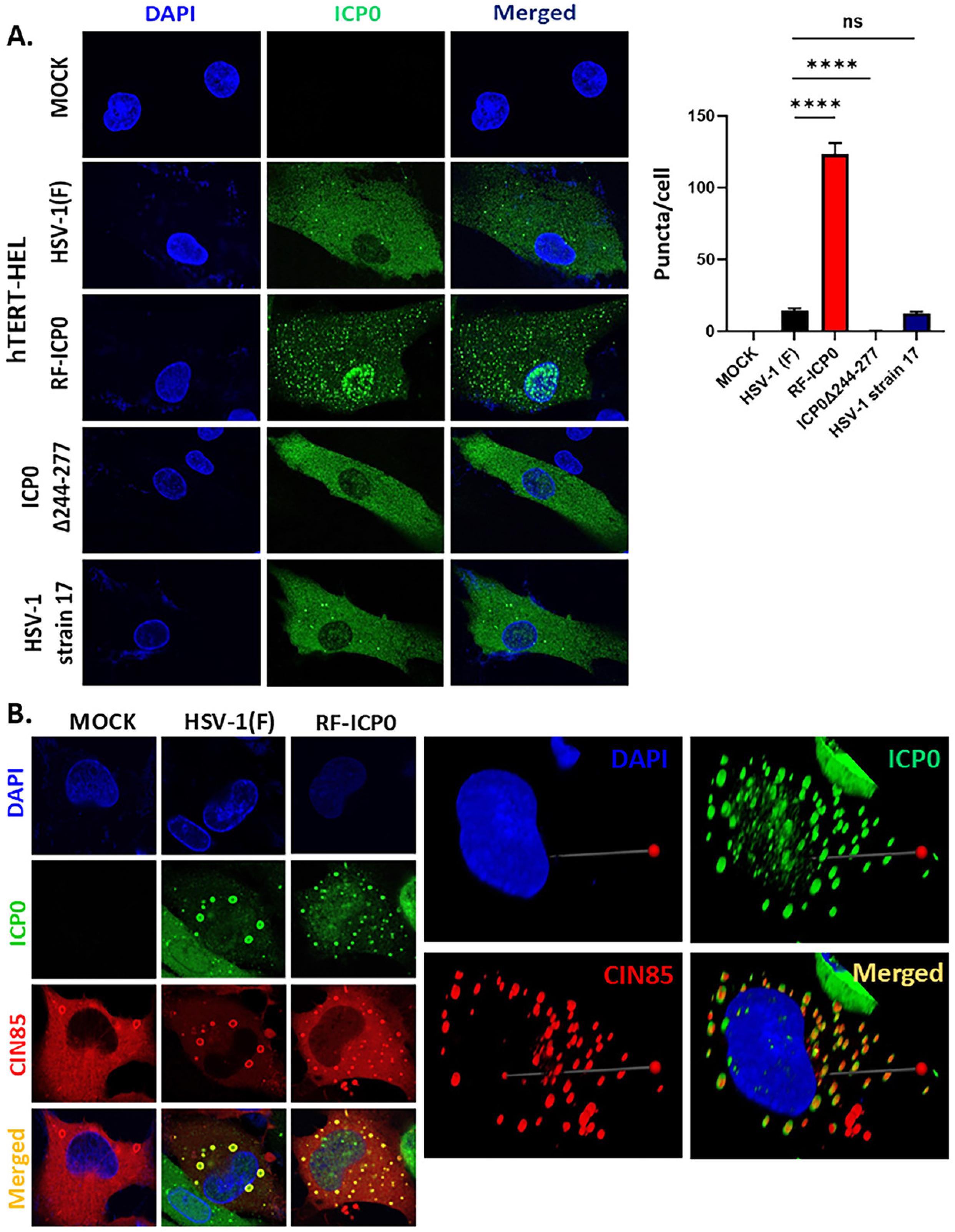
The RF domain of ICP0 affects CIN85 endosomes in hTERT-HEL and HEp-2 cells. **(A)** hTERT-HEL cells were infected with HSV-1(F), RF-ICP0, ICP0Δ244-277, and HSV-1 strain 17 (10 PFU/cell), or left uninfected. Cells were fixed at 15 h post-infection and stained with an ICP0 antibody. Cell nuclei were stained with DAPI. Images were captured using a Leica TCS SP8 confocal microscope. Puncta numbers per cell were quantified using Image J from approximately 45 cells from 3 independent experiments. Graphs were made using GraphPad Prism 10. **(B)** HEp-2 cells were transfected with a CIN85-Flag-expressing plasmid (250 ng/well). 24 h post transfection cells were infected with WT, RF-ICP0, and ICP0Δ244-277 (10 PFU/cell). Cells were fixed 16 h post infection and doubly reacted with ICP0 and CIN85 antibodies. Cell nuclei were stained using DAPI. Images were captured using a Leica TCS SP8 confocal microscope and analyzed using the Las X Software.

### CIN85 is required for ICP0 puncta formation, but not ATG5

We have previously demonstrated the importance of ICP0aa 244-277 for recruitment of ICP0 to CIN85 puncta using the ICP0Δ244-277 mutant virus. In a reciprocal approach we utilized a CIN85 KD hTERT-HEL cell line to determine the specificity of CIN85 as the recruiter of ICP0 to puncta sites. ATG5 is a known regulator of autophagic vesicles that we have previously shown to be present in CIN85/ICP0 vesicles (43), therefore we sought to determine if it had a role in ICP0 recruitment to vesicles using an ATG5 KD hTERT-HEL cell line. Cells were infected with WT, RF-ICP0, and ICP0Δ244-277 (5 PFU/cell) and fixed 15 h post-infection. Cells were stained with an ICP0 antibody and imaged using a Leica TCS SP8 confocal microscope (Figure 3A). Puncta number per cell were quantified using ImageJ (Figure 3B). We observed a decrease in puncta per cell in CIN85 KD hTERT-HEL cells in comparison to parental cells or ATG5 KD hTERT-HEL, indicating CIN85 to have a significant role in recruiting ICP0 to vesicular structures, but not ATG5. The efficiency of CIN85 and ATG5 depletion is depicted in Figure 3C. Overall, using both a mutant virus where ICP0 does not interact with CIN85 and a CIN85 depleted cell line we established that CIN85 is required for recruitment of ICP0 to CIN85 vesicles.

**Figure 3:**
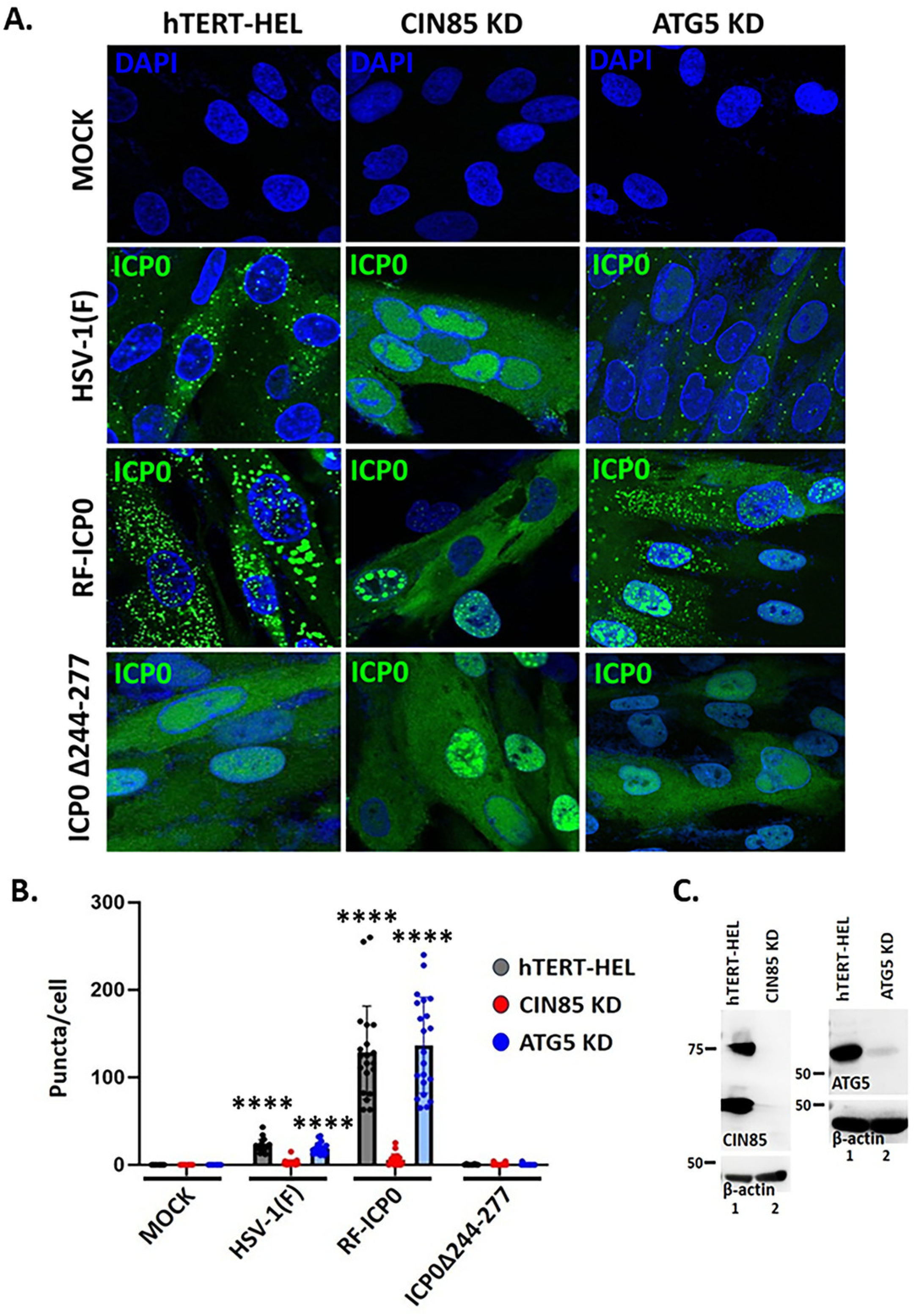
CIN85 is required for ICP0 recruitment to endosomes. **(A)** hTERT-HEL, hTERT-HEL CIN85KD, and hTERT-HEL ATG5KD cells were infected with HSV-1(F), RF-ICP0, ICP0Δ244-277 (5 PFU/cell) or left uninfected, and fixed at 15 h post-infection. Cells were stained for ICP0 and cell nuclei with DAPI. **(B)** Puncta numbers per cell were quantified using Image J from approximately 20 cells from 3 independent experiments. Graphs were made using GraphPad Prism 10. **(C)** Lysates were collected from hTERT-HEL, hTERT-HEL CIN85KD, and hTERT-HEL ATG5KD cells and analyzed by immunoblot to determine the efficiency of protein depletion.

### CIN85 and CD63 belong to distinct endosomes and populations of extracellular vesicles (EVs)

Our lab has extensively described a population of EVs released by HSV-1 infected cells that depend on the tetraspanin CD63 (56–58, 60–62). This protein typically traffics between the trans-Golgi network (TGN), plasma membrane, and endosomes and is highly enriched in a type of late endosomes called multivesicular bodies (MVBs) that traffic cargo to lysosomes for degradation or for exocytosis in EVs (63). CD63 sorts cargo to endosomes either via lateral associations with membrane-associated cargo through the formation of the tetraspanin web or through binding via the tetraspanin hypervariable loop (64–69). Previously we demonstrated that HSV-1 subverts the CD63 endosomal pathway to promote exocytosis of host antiviral factors, particularly membrane proteins, for immunoevasion (56, 58, 61). We also showed that CIN85 is exocytosed along with numerous host antiviral factors, particularly autophagosome components (43). Here, we sought to determine whether CD63 and CIN85 traffic through the same endosomal pathways and whether they are present in the same EV populations. To test this, we performed two sets of experiments. First, we analyzed the sub-cellular localization of CIN85 and CD63 during HSV-1 infection. For this, we transfected Hep-2 cells with a CD63-EGFP and a CIN85-Flag expressing plasmid for 24 h followed by infection with HSV-1(F) (10 PFU/cell). The cells were fixed at 16 h post-infection and stained with an ICP0 and a CIN85 antibody. As shown in Figure 4A, ICP0 and CIN85 colocalized in vesicular structures in the cytoplasm of infected cells, whereas CD63 displayed an ER-Golgi distribution and stained the nuclear envelope but did not colocalize with ICP0/CIN85. These data indicate that CIN85 and CD63 do not traffic through the same endosomal pathway. Second, to determine whether CIN85 and CD63 are packaged in the same or different EV populations we analyzed the EVs released by HSV-1(F) infected cells. For this, hTERT-HEL cells were infected with the WT virus (0.3 PFU/cell) and at 17 h post-infection culture supernatant was collected, clarified at low-speed centrifugation to remove floating cells and larger debris, filtered/concentrated through 100 kDa cutoff filters, and subjected to ultracentrifugation through a 6-18% iodixanol/sucrose gradient, as we have described before (56–58). Fractions of 500 µl were collected from the top to the bottom of the gradient and analyzed for CIN85, CD63, and the HSV-1 capsid protein VP5 to indicate virus presence. As shown in Figure 4B, CIN85 and CD63 accumulate in different fractions of this gradient with CIN85+ EVs being lighter than the CD63+ EVs. Following their exocytosis, intracellular CIN85 and CD63 display reduced levels (Figure 4B).

**Figure 4:**
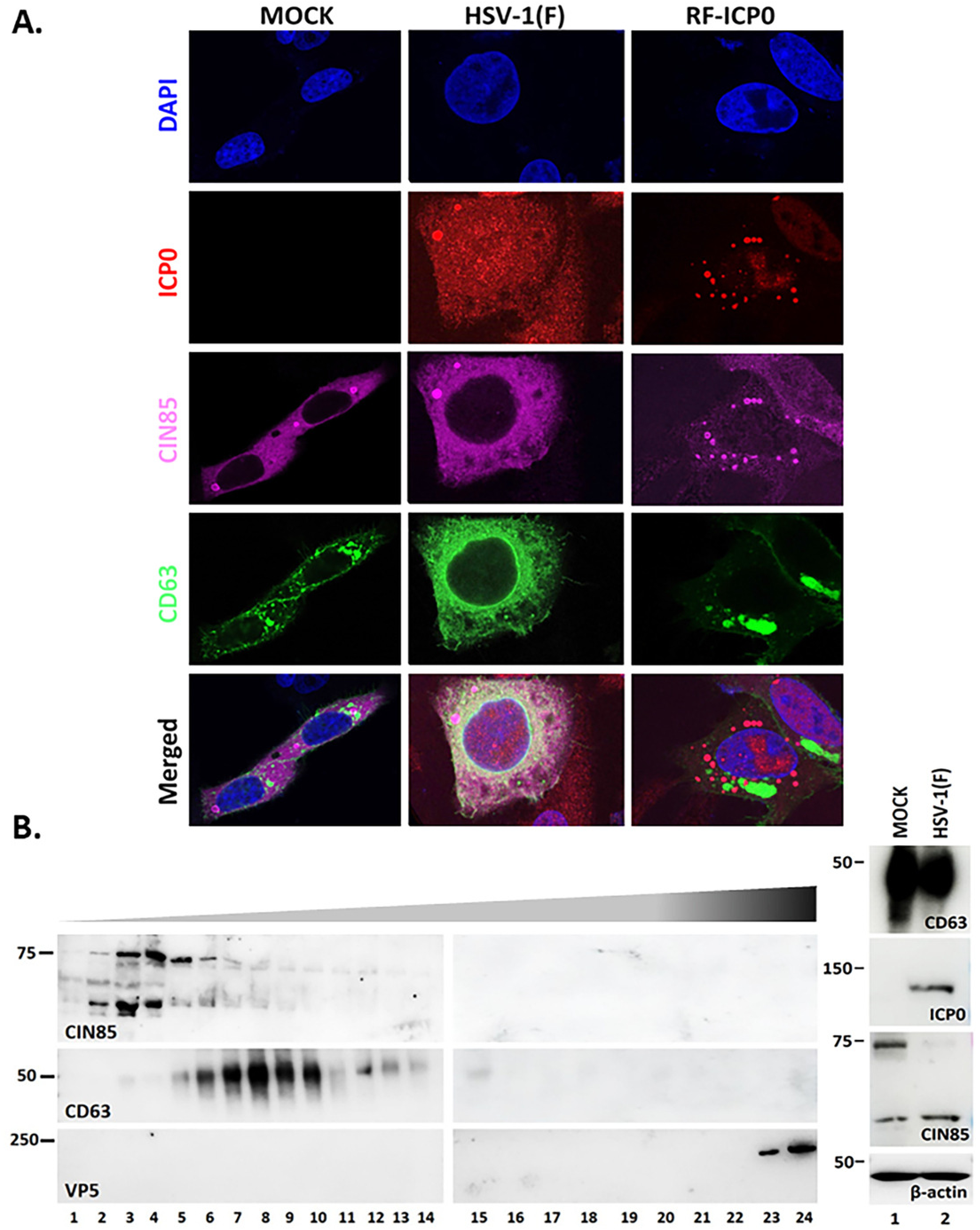
CIN85 and CD63 belong to distinct endosomal and exocytosis pathways. **(A)** HEp-2 cells were transfected with CIN85-Flag (250 ng/well) and an EGFP-CD63 (250 ng/well) expressing plasmid. Cells were infected at 24 h post-transfection with WT, RF-ICP0 (10 PFU/cell), or left uninfected. Cells were fixed 16 h post-infection and doubly stained using ICP0 and CIN85 antibodies. Alexa Fluor 594 nm and Alexa Flour 680 nm antibodies were used to visualize ICP0 and CIN85 respectively. Images were captured using a Leica TCS SP8 confocal microscope. **(B)** hTERT-HEL cells were uninfected or infected with WT virus (0.3 PFU/cell) and supernatant was collected 17 h post-infection. Supernatant was centrifuged, filtered, and concentrated using Centricon Plus 70 (100-kDa cutoff) to 1 mL. Sample was loaded on top of an iodixanol/sucrose gradient. Gradient preparation is further detailed in the Materials and Methods. Fractions (500 ul) were collected from the top to the bottom of the gradient, totaling 24 fractions. Equal volume of the fractions was analyzed via immunoblot for CD63, CIN85, and the viral protein VP5. Equal amounts of proteins from total cell lysates were analyzed for CD63, CIN85, ICP0, and β-actin.

Taken together, these studies indicate that CIN85 and CD63 belong to distinct endosomes and population of EVs and ICP0 selectively colocalizes with CIN85.

### ICP0/CIN85 vesicles contain autophagy markers, but they are not autophagosomes and they do not colocalize with lysosomes

Endocytic vesicles can either recycle cargo to the plasma membrane, distribute cargo to other membrane compartments inside the cells, target cargo to lysosomes for degradation or divert it for exocytosis (70–75). Lysosomal degradation may or may not involve the autophagy pathway (74–78). Autophagy is the homeostatic process for cell protein and organelle degradation. There are several types of autophagy, including macroautophagy which functions to seclude and degrade cytoplasmic cargo via the formation of cytosolic vesicles called autophagosomes. Autophagosomes typically fuse with lysosomes upon which a variety of enzymatic activities within lysosomes will degrade autophagic cargo (79–81).

We sought to determine if the ICP0/CIN85 vesicles fuse with autophagosomes or with lysosomes to degrade their cargo. We first determined if the ICP0/CIN85 vesicles colocalize with autophagosome markers such as LC3 (microtubule-associated protein 1A/1B-light chain 3) and ATG5 (autophagy-related protein 5) that drive autophagosome formation, or with p62/SQSTM1 (sequestosome 1), an autophagy adaptor protein. HEp-2 cells were co-transfected with CIN85-Flag and either mCherry-LC3, mCherry-ATG5, or mCherry-p62 plasmids. At 24 h post-transfection, the cells were infected with either the WT virus or RF-ICP0 (10 PFU/cell). The cells were fixed at 16 h post-infection and images were acquired with the same settings of a Leica TCS SP8 confocal microscope. As shown in Figure 5A, LC3, ATG5, and p62/SQSTM1 colocalized with ICP0 and CIN85 in vesicular structures. These data indicate that autophagosome markers are recruited to ICP0/CIN85 vesicles and that this recruitment is not dependent upon the ICP0 E3 ubiquitin ligase activity.

**Figure 5:**
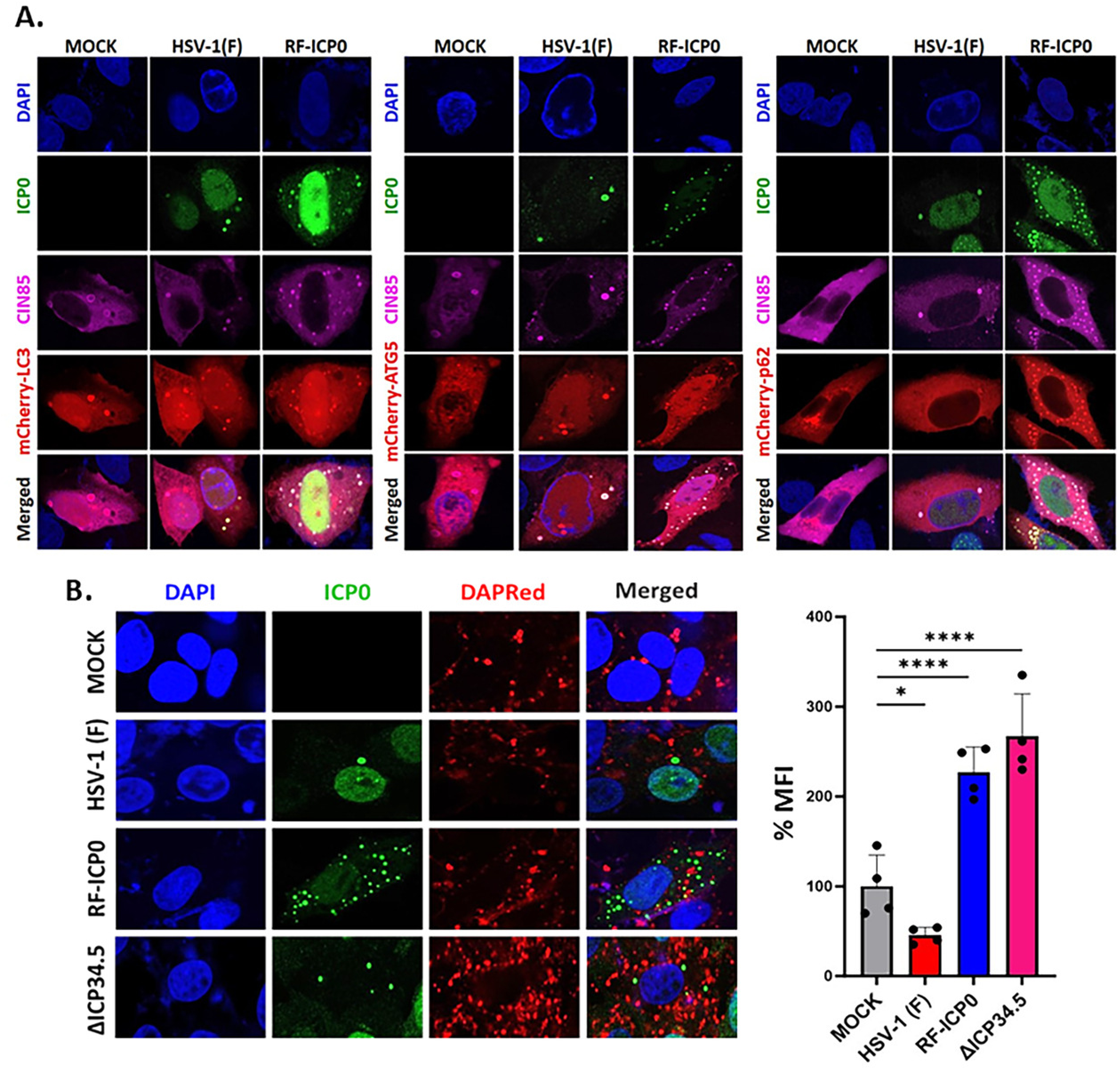
ICP0/CIN85 endosomes do not fuse with autophagosomes. **(A)** HEp-2 cells were transfected with a CIN85-Flag (250 ng/well) and an mCherry-LC3, mCherry-p62, or mCherry-ATG5 plasmid (250 ng/well). Cells were infected 24 h post-transfection with WT virus, RF-ICP0 (10 PFU/cell), or left uninfected. Cells were fixed at 16 h post-infection and doubly stained for ICP0 and CIN85. Images were captured using a Leica TCS SP8 confocal microscope. **(B)** HEp-2 cells were transfected with a CIN85-Flag-expressing plasmid (250 ng/well). Cells were infected 24 h post transfection with WT virus, RF-ICP0, or left uninfected (10 PFU/cell). Cells were washed with 1% DMEM, then incubated with DAPRed (0.1 µmol) in 1% DMEM at 37 ^0^C for 30 min. Cells were fixed 4 h later and stained for ICP0. Cell nuclei were stained using DAPI. DAPRed mean fluorescence intensity (MFI) was calculated using the average of four independent experiments with ImageJ. The graph was created using GraphPad Prism 10.

To determine whether ICP0/CIN85 vesicle fuse with autophagosomes we used DAPRed, a small molecule that incorporates at the sites of double membrane elongation and therefore serves an autophagosome indicator by emitting light in hydrophobic conditions. We transfected Vero cells with a CIN85-Flag expressing plasmid for 24 h followed by infection with WT virus (10 PFU/cell). Live cells were stained with DAPRed 12 h post-infection for 30 minutes. The cells were then washed, fixed 4 h post-treatment and stained with an ICP0 antibody. We observed no colocalization between ICP0 and DAPRed, indicating no autophagosome fusion with ICP0 vesicles (Figure 5B). We also monitored the ICP0 vesicles after infection with RF-ICP0 or ΔICP34.5 viruses (10 PFU/cell). Both mutant viruses display defects in evading autophagy effectively; therefore, we observed an increase in DAPRed fluorescence following signal quantification indicative of increased autophagosome formation (Figure 5B). However, no colocalization of ICP0 vesicles with autophagosomes was observed.

Endosomes can also fuse to lysosomes directly without involving autophagy (74–76, 78). To investigate this possibility for the ICP0/CIN85 vesicles we tested whether they colocalize with LAMP1 (lysosomal associated membrane protein 1). LAMP1 and LAMP2 comprise 50% of all lysosomal proteins and are thought to have an integral role in maintaining lysosomal integrity, pH, and catabolism (82–84). We transfected HEp-2 cells with a LAMP1-RFP and CIN85-Flag expressing plasmid for 24 h followed by infection with WT virus (10 PFU/cell). Cells were fixed 16 h post-infection and stained with an ICP0 and CIN85 antibody. We observed no colocalization between LAMP1 and ICP0/CIN85 vesicles (Figure 6A). We also tested colocalization of LAMP1 with ICP0/CIN85 vesicles in RF-ICP0 and ΔICP34.5-infeced cells (10 PFU/cell) and no colocalization was observed (Figure 6A).

**Figure 6:**
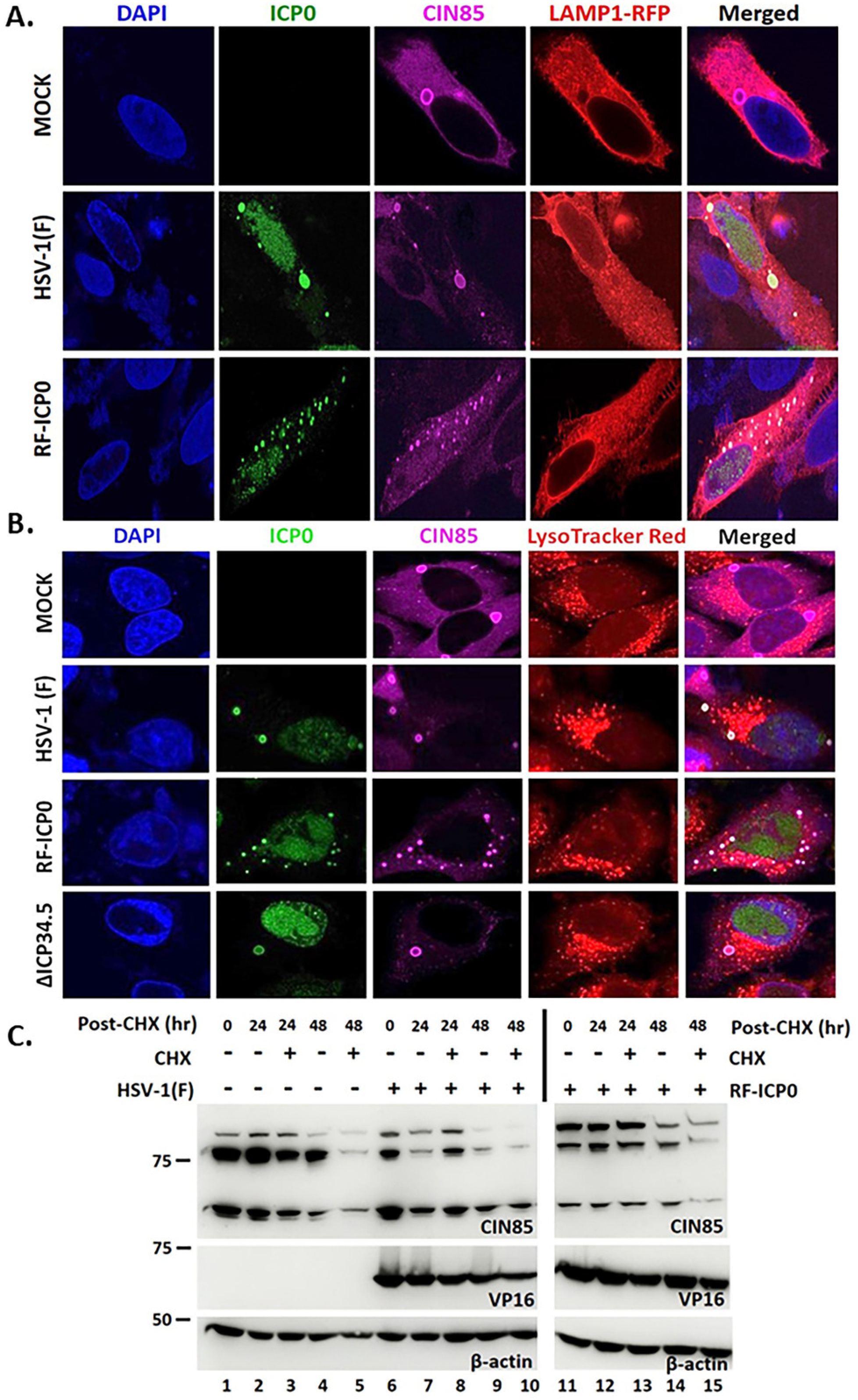
ICP0/CIN85 endosomes do not fuse with lysosomes. **(A)** HEp-2 cells were transfected with a CIN85-Flag (250 ng/well) and a LAMP1-RFP (250 ng/well) plasmid. Cells were infected 24 h post-transfection with WT virus and RF-ICP0 (10 PFU/cell) or left uninfected. Cells were fixed 16 h post-infection and doubly stained for ICP0 and CIN85. Images were captured as before. **(B)** HEp-2 cells were transfected with a CIN85-Flag-expressing plasmid (250 ng/well). Cells were infected 24 h post-transfection with WT virus, RF-ICP0, ΔICP34.5 (10 PFU/cell), or left uninfected. Cells were washed with 1% DMEM, then incubated with LysoTracker Red (0.1 µmol) in 1% DMEM at 37^0^C for 30 min. Cells were fixed 4 h post-treatment and stained for ICP0. Cell nuclei were stained using DAPI. Images were captured as before. **(C)** hTERT-HEL cells were infected with WT virus, RF-ICP0 (5 PFU/cell) or remained uninfected. 12 h post infection cells were treated with CHX (100 µg/μl), and cell lysates were collected 0, 24, and 48 h post CHX treatment. Equal amounts of proteins from total cell lysates were analyzed for CIN85. VP16 served as a control for the infection and β-actin as a loading control.

In a complementary approach, we determined whether ICP0/CIN85 vesicles colocalize with lysosomes stained with LysoTracker Red. Staining with this dye utilizes a protonation-based trapping strategy in which under cytosolic pH conditions the molecule remains largely unprotonated, allowing for free diffusion across biological membranes. Upon reaching compartments with lower pH, such as lysosomes, the molecule becomes charged and can no longer freely diffuse across membranes, leading to accumulation within the compartments. We transfected Vero cells with a CIN85-Flag expressing plasmid for 24 h followed by infection with WT virus (10 PFU/cell). Live cells were stained with LysoTracker Red 12 h post-infection for 1 h. Then, cells were washed, fixed 2 h post treatment and stained with an ICP0 and CIN85 antibody. We observed no colocalization between ICP0 and LysoTracker Red, indicating no fusion of ICP0/CIN85 vesicles with lysosomes (Figure 6B). We performed similar analysis in RF-ICP0 and ΔICP34.5-infeced cells (10 PFU/cell) and no colocalization of ICP0/CIN85 vesicles with LysoTracker Red was observed (Figure 6B). Since in RF-ICP0-infected cells CIN85 is not exocytosed we asked if degradation occurs intracellularly. For this, we treated the infected cells (5 PFU/cell) with cycloheximide (CHX) at 12 h post-infection or left them untreated, harvested them at 0, 24, and 48 h post-CHX treatment and analyzed them for CIN85. We determined that the half-life of CIN85 in uninfected cells is longer than 24 h, however in RF-ICP0-infected cells the stability of CIN85 decreases. These data indicate that CIN85 is degraded to some extent in RF-ICP0-infected cells.

Also, we conducted a lysosome activity assay to determine the effect of HSV-1 infection, particularly through the role of ICP0 in endocytosis. DQ BSA (dye-quenched bovine serum albumin), a self-quenching dye conjugated to bovine serum albumin, releases fluorescent signal upon digestion by hydrolases within the lysosome, therefore providing insight on lysosomal activity. We infected HEp-2 cells with WT, RF-ICP0, and ICP0Δ244-277 (10 PFU/cell), then at 4 h and 12 h post-infection cells were incubated with DQ-BSA (10 µM) for 30 minutes in 37 ^0^C. Cells were then washed, and live cell imaging was performed 4 h later. Rapamycin, a known mTOR (mechanistic target of rapamycin) inhibitor that induces autophagy, was used as a positive control, by pretreating cells (5 µM) for 2 h prior to DQ BSA treatment. We observed an increase in DQ-BSA fluorescence signal at 8 h post-infection, with WT virus inducing almost 2 times the fluorescence of uninfected cells (Figure 7A-B). DQ-BSA signal was also induced by the ICP0 mutant viruses but to lower extent than WT virus (Figure 7A-B). The DQ-BSA fluorescent signal was significantly reduced by 16 h post-infection by all viruses (Figure 7A-B).

**Figure 7:**
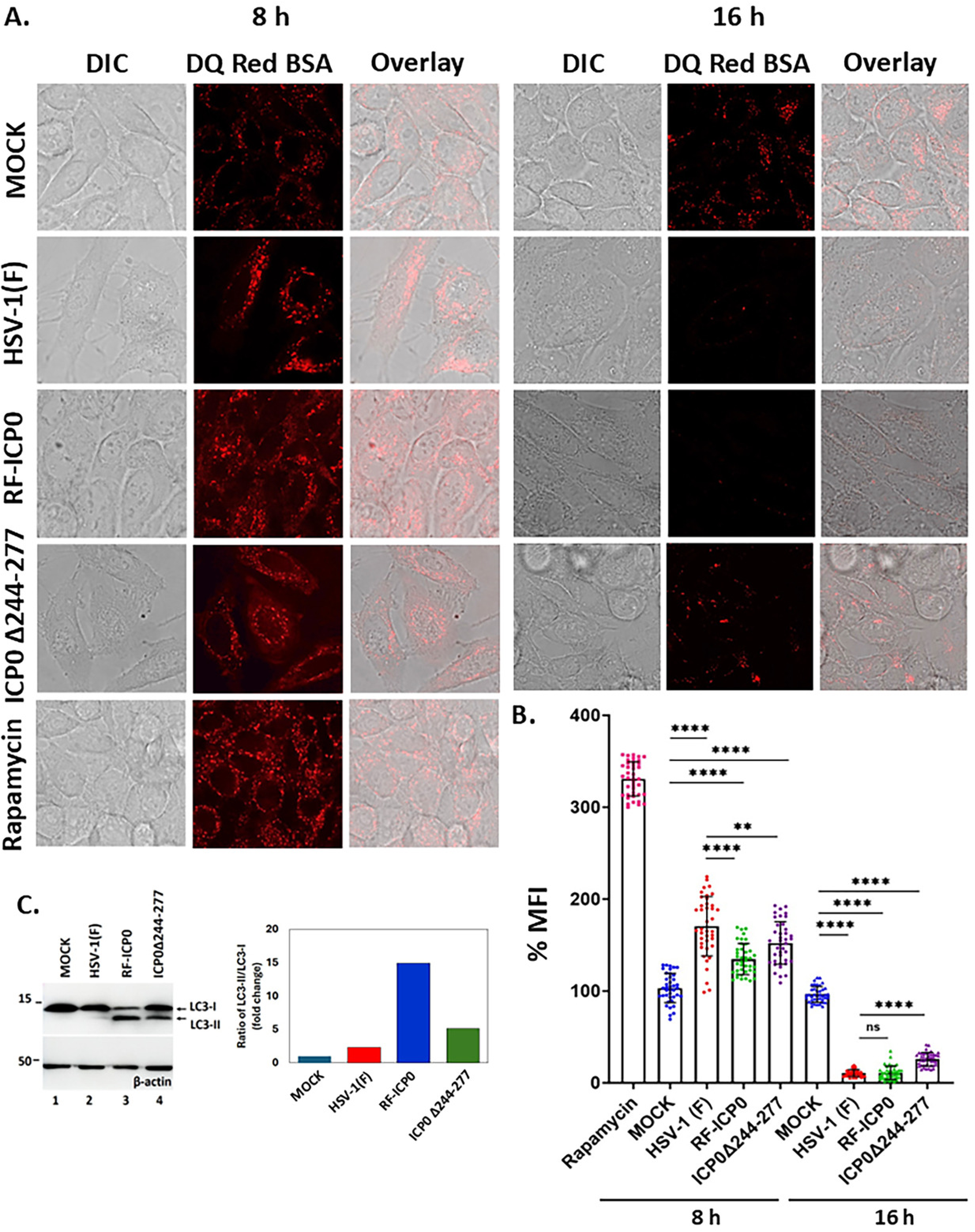
Lysosomal activity during HSV-1 Infection. **(A)** HEp-2 cells were infected with WT, RF-ICP0, and ICP0Δ244-277 (10 PFU/cell) or left uninfected for either 4 or 12 h at which time DQ Red BSA was added for 30 min at 37 ^0^C. Cells were live imaged 4 h post DQ Red BSA staining. As a positive control, rapamycin cells were pretreated with 5 µM rapamycin for 2 h and live imaged 4 h post DQ Red BSA staining. Images were captured using a Leica TCS SP8 confocal microscope. **(B)** MFI was calculated using 40 cells from three independent experiments with ImageJ. The graph was created using GraphPad Prism 10. **(C)** hTERT-HEL cells were infected with WT virus, RF-ICP0, ICP0Δ244-277 (5 PFU/cell) or left uninfected. Cells were harvested at 9 h post-infection and equal amounts of proteins from total cell lysates were analyzed for LC3 lipidation with β-actin serving as a loading control. The LC3 signals were quantified using ImageJ and normalized to β-actin. LC3-II/LC3-I ratio for each sample relative to uninfected cells is depicted.

Finally, we analyzed the lipidation of LC3 as an indication of autophagosome formation after infection with WT virus and the ICP0 mutants. The analysis was performed at 9 h post-infection (5 PFU/cell), where we observed that RF-ICP0 infection causes extensive LC3 lipidation (Figure 7C). The LC3 lipidation was less extensive by ICP0Δ244-277 infection, whereas only minimal LC3 lipidation was observed with WT virus (Figure 7C). These results were expected as WT virus has evolved elegant strategies to evade autophagy, whereas the ICP0 mutants lack certain ICP0 functions required for evasion of antiviral signaling (17, 23, 29, 85–89).

Overall, these data indicate that the ICP0/CIN85 vesicles are distinct from autophagosomes and they do not fuse with lysosomes. Also, we observed an initial increase in lysosomal activity followed by a reduction later during infection, which coincides with an elevation in LC3 lipidation found in the ICP0 mutant viruses.

### Exocytosis of ICP0/CIN85 cargo requires both binding of ICP0 to CIN85 and intact E3 ubiquitin ligase activity

We have previously reported that HSV-1 infected cells produce about 2-fold more EVs compared to uninfected cells that carry virus-specific cargo and display an overall antiviral effect on recipient cells (56–58, 60–62). Here, we sought to determine the effect of the ICP0 E3 ubiquitin ligase activity and ICP0/CIN85 interaction on the biogenesis and cargo of these EVs, and their overall antiviral effect. For this, hTERT-HEL cells were left uninfected or infected with WT, RF-ICP0, and ICP0Δ44-277 viruses (0.5 PFU/cell). Culture supernatant was collected at 48 h post-infection, total EVs were pelleted, as detailed in Materials and Methods, and analyzed for markers of EVs and cargo we previously found to be present in ICP0/CIN85 vesicles (Figure 8A) (43). We found that both CIN85 and EGFR, a receptor known to be endocytosed by CIN85, were exocytosed in HSV-1-infected cells and both the binding of ICP0 to CIN85 and the ICP0 E3 ubiquitin ligase activity were required for this exocytosis. Also, Rab5, Rab7, LC3-II, and p62/SQSTM1 that we previously reported in ICP0/CIN85 vesicles were exocytosed in an ICP0-dependent manner. Factors that are known to be targeted by ICP0 for proteasomal degradation such as SP100A and its sumoylated forms were also exocytosed in an ICP0-dependent manner. Galectin 3, a member of the β-galactoside-binding proteins, with roles in host defense, microglia activation, cell proliferation, and apoptotic regulation (90–94) was exocytosed in an ICP0-dependent manner. However, not all infection-dependent protein exocytosis requires ICP0-RF or CIN85 interaction. For example, LAMP1, the lysosomal marker we previously observed to not colocalize with ICP0/CIN85 vesicles, was found to be exocytosed during infection in an ICP0-RF- and CIN85-independent manner. CD63 exocytosis was not significantly impacted by the ICP0 mutants.

**Figure 8:**
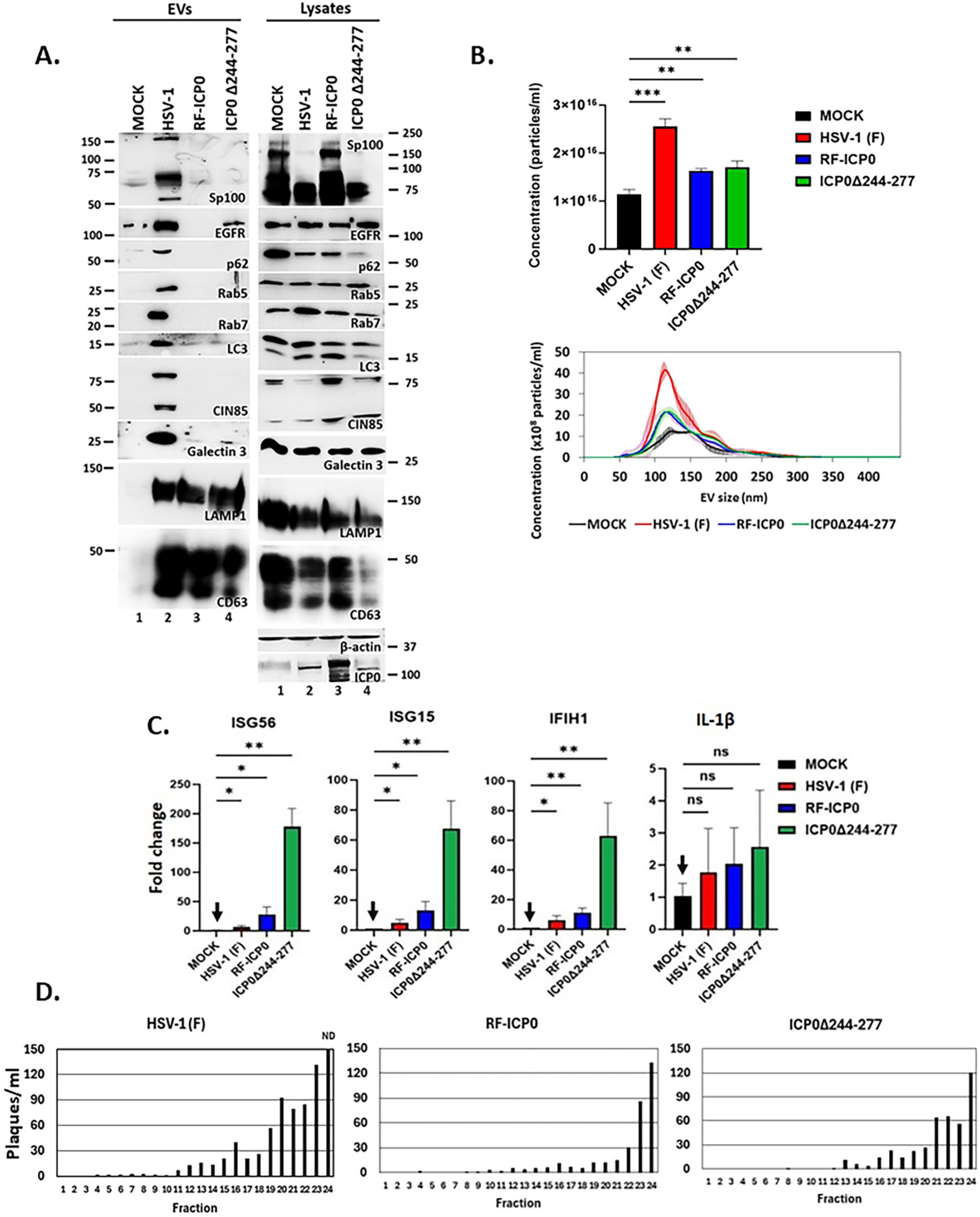
Intact ICP0 E3 ubiquitin ligase activity and binding of ICP0 to CIN85 are required for exocytosis during HSV-1 infection. **(A)** hTERT-HEL cells were infected with WT virus, RF-ICP0, ICP0Δ244-277 or left uninfected (0.5 PFU/cell), and culture supernatant was collected at 48 h post-infection. EVs were isolated and pelleted as detailed in Materials and Methods. Equal amounts of proteins from EVs and cell lysates were analyzed by immunoblot. **(B)** hTERT-HEL cells were infected with WT virus, RF-ICP0, ICP0Δ244-277 (0.5 PFU/cell), or remained uninfected and supernatant was collected 17 h post-infection as detailed in Materials and Methods. EVs were loaded onto an iodixanol-sucrose gradient (6-18%), centrifuged in a SW41Ti rotor for 150 min at 200,000x *g* and 4 ^0^C, and 500 μl fractions were collected from the top to the bottom of the gradient (total 24 fractions). Fractions 1-8 were pooled and analyzed by NTA. The quantities of the EVs and their size distribution are depicted from 3 independent EV isolations. The color-shaded regions represent the error bars showing the variation in concentration and size of EVs between different conditions. The average total EV concentrations between samples were plotted using GraphPad Prism 10. **(C)** EVs were exposed to uninfected hTERT-HEL cells. EVs from Mock cells were used at 1000 EVs/cell, while EVs/cell from infected samples were normalized to Mock based on the average total number of EVs produced during each condition. 12 h post exposure total RNA was extracted and analyzed by RT-qPCR. **(D)** EV purity was verified by plaque assays. All experiments were done in triplicate.

Next, we analyzed the biophysical properties of EVs released after infecting cells with the ICP0 mutant viruses and their role in antiviral immunity. hTERT-HEL cells were infected with WT, RF-ICP0, and ICP0Δ244-277 (0.3 PFU/cell) or left uninfected. Culture supernatants were collected at 17 h post-infection, filtered, and concentrated. EVs were loaded onto an iodixanol-sucrose gradient (6-18%) and 500 μl fractions were collected from the top to the bottom of the gradient (total 24 fractions). The fractions were analyzed for EV markers by Western blot and for purity by plaque assays to ensure separation of EVs from virus (Figure 8D). The fractions containing the EV markers (top 8) were pooled together, analyzed using nanoparticle tracking analysis (NTA), and further washed and concentrated. We determined that there is a reduction in EV concentration, though size remained similar, when comparing WT to both ICP0 mutant viruses (Figure 8B). We then utilized 1,000 EVs/cell from uninfected cells, which corresponds to number of EVs present in the supernatant of uninfected cultures within 24 h, and normalized the dose of EVs from infected cells to best mimic natural infection conditions. EVs were exposed to uninfected hTERT-HEL cells for 12 h and total RNA was extracted and analyzed via RT-qPCR. We observed an increase in innate immune reactions of recipient cells following exposure to EVs from both mutant viruses in comparison to EVs from WT virus (Figure 8C). Activation of innate immunity was negligible with EVs from uninfected cells (Figure 8C).

These findings indicate that ICP0 is recruited by CIN85 on endosomes where through its E3 ubiquitin ligase activity determines the CIN85 endosome and EV proteome content. These EVs shape the microenvironment of infection by activating antiviral responses.

## Discussion

ICP0 is critical for productive HSV-1 infection and virus reactivation from latency, thereby it has been extensively studied since its discovery more than six decades ago (12, 16, 31, 95–98). ICP0 is known as the promiscuous transactivator of the virus needed for viral gene expression (3, 7, 8, 99). In addition, through its E3 ubiquitin ligase activity, it mediates the degradation of cellular substrates such as constituents of the ND10 bodies and other innate immunity components (13, 18, 31, 50, 55, 99–101). Most known ICP0 functions are performed in the nucleus where the protein initially localizes; however, after replication initiates, ICP0 accumulates in the cytoplasm where its further functions are unknown. We recently reported that ICP0 interacts and colocalizes with CIN85 on the surface of vesicular structures in the cytoplasm of infected cells (43). Here, we sought to determine the potential role of the ICP0 E3 ubiquitin ligase in the formation and function of ICP0/CIN85 endosomes. The salient findings of our studies can be summarized as follows:

We discovered that disruption of the ICP0 RF domain is associated with an endosomal phenotype, characterized by a larger number of smaller ICP0/CIN85 vesicles than WT virus infections. This phenotype is independent of cell type and is observed both in human primary and immortalized cell cultures. Disruption of the RF domain abrogates ICP0’s E3 ubiquitin ligase activity, thus, our findings indicate that ICP0, through its catalytic activity, likely modifies CIN85 endosomes. This modification could influence the biogenesis and formation of CIN85 endosomes, cargo, maturation, trafficking, or their ability to fuse with target compartments. Consistently, we observed that in RF-ICP0 infections CIN85 endosomes cannot be exocytosed like in WT virus infections. CIN85 is typically involved in degradative or recycling pathways, this is why only trace amounts of CIN85 were detected in EVs from uninfected cells. This indicates that the WT virus encodes a function that diverts the CIN85 endosomes for exocytosis. Our studies have linked this function to ICP0, and it is performed on endosomes while requiring intact E3 ubiquitin ligase activity, as neither RF-ICP0 nor ICP0Δ244-277 viruses promote CIN85 exocytosis. An attractive hypothesis is that ICP0 facilitates cargo flow through the CIN85 pathway, whereas catalytically inactive ICP0 reduces cargo efflux and prevents CIN85 endosomes from coalescing into a few larger endosomes. Another interpretation of this phenotype is the buildup of ICP0 during RF-ICP0 virus infection, due to the absence of autocatalytic degradation, might form a physical block preventing vesicle trafficking and fusion. Nevertheless, this is the first study linking ICP0 catalytic activity with endosomal activity.

Another discovery we made is that the CIN85+ EVs are distinct from the CD63+ EVs we have previously described and they originate through different endosomal pathways (56–58, 61). CD63 typically traffics between plasma membrane, TGN, late endosomes, lysosomes, and EVs (63, 102–104). In HSV-1-infected cells, CD63 endosomes are diverted for exocytosis and carry with them antiviral factors (56–58, 61). CIN85 is an endocytic adaptor found in tubular endosomes, recycling endosomes, and the degradative pathway, which regulates cell surface receptors (44–46, 48). CIN85 is required for the recruitment of ICP0 to CIN85 endosomes, whereas other vesicular regulators such as ATG5, an autophagic vesicles regulator, have no role. In CIN85 depleted cells or after infection with ICP0Δ244-277, where ICP0 does not interact with CIN85, ICP0 remains diffuse in the cytoplasm. Through this interaction, the virus likely remodels the cell surfaceoma and diverts selected factors for exocytosis. Failure of ICP0 to interact with CIN85 results in decreased progeny virus production due to elevated antiviral responses (43).

An additional discovery we made is related to the crosstalk between the CIN85 endocytic pathway and the autophagolysosomal pathway. The CIN85 endosomes colocalize with several autophagosome components, such as ATG5, LC3, and p62/SQSTM1, both in WT virus-infected and RF-ICP0-infected cells (43). Through the CIN85 pathway autophagosome and lysosome components are extruded from infected cells and this likely represents a complementary strategy through which HSV-1 evades autophagy (43). However, CIN85 endosomes were not entering the autophagolysosomal pathway, even when autophagy was activated, such as after infection with RF-ICP0 or ΔICP34.5 viruses (85, 87). With these mutant viruses we observed an increase in LC3 lipidation, a hallmark for autophagosome formation, an increase in the number of autophagosomes, abundant lysosomes, but the ICP0/CIN85 endosomes did not colocalize with autophagolysosomes. This may contribute to the increased number of ICP0/CIN85 endosomes in RF-ICP0 infected cells, as they are neither degraded nor exocytosed. Further, we found that CIN85 has a long half-life, over 24 h, which could further contribute to the RF-ICP0 phenotype. However, we did observe a decline in CIN85 levels in RF-ICP0-infected cells that occurred between 24 and 48 h post-CHX treatment. This decline could occur through a lysosomal pathway, proteasomes, or other degradation mechanisms. Notably, in HSV-1 productively infected cells lysosomal activity increases during the early stages of infection, followed by a decline at later stages. This was the case with ICP0 mutant viruses as well. The decrease in lysosomal activity during HSV-1 infection has been reported before and is mostly associated with defects in trafficking of lysosomal enzymes (105, 106). In our assays we used DQ Red BSA, which releases fluorescent signal upon digestion of hydrolases found within the lysosome, and observed a marked decrease in the fluorescent signal during the late stages of infection, supporting defects in the lysosomal trafficking pathway.

A final discovery has to do with the impact of EVs produced by infected cells in autologous uninfected cells. RF-ICP0- or ICP0Δ244-277-infected cells release less EVs compared to WT virus-infected cells that do not carry the CIN85-associated antiviral factors we have reported before (43, 58, 60). However, these EVs are more potent at activating type I-IFN and pro-inflammatory responses. Thus, future studies aim at identifying this cargo.

Overall, we have provided evidence that the ICP0 E3 ubiquitin ligase activity plays a role in the CIN85 endosomal pathway and is required for exocytosis of selected antiviral factors for immunoevasion. These findings further highlight that the endocytic and exocytic pathways are integral part of HSV-1 life cycle; they are subverted by the virus for immunoevasion and they shape the microenvironment of infection likely driving pathogenesis.

## Materials and Methods

### Cell lines, viruses, and chemicals

Adult primary human epidermal keratinocytes (HEKa), Vero, HEp-2, ARPE-19, 293T, and U2OS cells were obtained through ATCC and cultured in accordance with manufacturer’s instructions. HaCaT were obtained through Abm and cultured according to supplier’s instruction. hTERT-HEL cells (immortalized human embryonic lung fibroblasts, human telomerase reverse transcriptase [hTERT] transformed) were a gift from Dr. Roizman B (University of Chicago). HSV-1 strain F and HSV-1 strain 17 have been described before (107, 108). The properties of RF-ICP0, ICP0Δ244-277 and ΔICP34.5 viruses were described before (43, 59, 109).

### Development of stable cell lines and transfection assays

Mission pLKO.1 based plasmids that carry shRNAs for the depletion of human CIN85 and ATG5 were purchased from Sigma. For the development of the respective lentiviral vectors, 293T cells seeded in a F25 cm2 flask at a 60% confluency were transfected with 2.3 μg of the plasmid carrying the shRNA, 2.3 μg of the Gag-Pol–expressing plasmid, and 300 ng of the VSV-G–encoding plasmid with Lipofectamine 3000, according to manufacturer’s instruction (Thermo Fisher Scientific). At 48 h after transfection, the supernatant from the cultures was collected, filtered through 45-μΜ pore-size filters, and used to infect hTERT-HEL cells in the presence of polybrene (10 μg/ml). Puromycin selection (2 μg/mL) was initiated 24 h after exposure to lentiviruses and continued until only resistant clones emerged. The clones of hTERT-HEL cells, with the greater depletion in protein of interest, were used for those studies. The stable lines were maintained in the presence of puromycin, which was removed 24 h before assays were performed. Transfection assays were calculated with the ratio of plasmid to Lipofectamine 3000 for a given number of cells in accordance with the manufacturer’s instructions. Several plasmids used in this study were purchased through Addgene: pmCherry-ATG5 (#13095), mCherry-hLC3B-pcDNA3.1 (#40827), LAMP1-RFP-3Xha-pLJC5 (#102932), mCherry-Sequestosome1-N1 (#55132). The pcDNA-CIN85-Flag was gifted by Ivan Dikic (Goethe University Frankfurt).

### Immunoblot analysis

Cells were lysed in triple detergent buffer (50 mM Tris-HCl [pH 8], 150 mM NaCl, 0.1% sodium dodecyl sulfate, 1% Nonidet P-40, 0.5% sodium deoxycholate, 100 mg/ml of phenylmethylsulfonyl fluoride) supplemented with phosphatase inhibitors (10 mM NaF, 10 mM b-glycerophosphate, 0.1 mM sodium vanadate) and protease inhibitor cocktail (Sigma). Samples were briefly sonicated and protein concentration determined using the Bradford method (Bio-Rad Laboratories). Proteins were visualized with ECL western blotting detection reagents (Pierce). Primary antibodies used at 1:1000 dilution from Santa Cruz: ICP0 (sc-53070), CD63 (sc-5275), Rab5 (sc-46692), ATG5 (sc-133158), Rab7 (sc-376362), EGFR (sc-373746), CIN85 (sc-166862), β-actin (sc-47778), galectin 3 (sc-32790), and Lamp1 (sc-20011). LC3B rabbit polyclonal antibody (NOVUS Biologicals; NB100-2220) was used at 1:1000. CIN85 rabbit (Cell Signaling #12304) and p62/SQSTM1 mouse (Cell Signaling #88588) monoclonal antibodies were used at 1:1000. Sp100 rabbit polyclonal antibody (GeneTex #GTX131569) was used at 1:1000.

### Immunofluorescence analysis

The procedures were described elsewhere (43). Briefly, the cells were fixed in 4% paraformaldehyde, permeabilized, blocked with PBS–TBH solution consisting of 0.1% Triton X-100 in PBS, 10% horse serum, and 1% BSA, and reacted with primary antibodies (listed above) diluted in PBS–TBH. The cultures were rinsed with PBS–TBH and reacted with appropriate Alexa-Fluor-conjugated secondary antibodies, diluted 1:1,000 in PBS–TBH. Finally, cells were washed with PBS–TBH and the samples were mounted in VECTASHIELD Antifade Mounting Medium with DAPI (H-1200) and imaged using a Leica TCS SP8 confocal microscope. Alexa-Flour-conjugated antibodies were used at 1:1000 dilution and obtained from Invitrogen: Alexa Fluor 594 goat anti-rabbit IgG (A11012), Alexa Fluor 594 goat anti-mouse IgG (A11005), Alexa Fluor 488 goat anti-rabbit IgG (A11008), Alexa Fluor 488 goat anti-mouse IgG (A11001), Alexa Fluor 680 goat anti-rabbit IgG (A21109).

### DAPRed autophagy stain assay

Vero cells were seeded at 70–80% confluency on 4-well slides. Cells were transfected with a CIN85-Flag plasmid using Lipofectamine 3000 Transfection kit (Thermo Fisher Scientific) and then 24 h later were infected with WT, RF-ICP0, or ΔICP34.5 (10 PFU/cell). DAPRed stain (Dojindo Molecular technologies Inc. NC1879567) was added 12 h post-infection for 30 min at 37 ^0^C (0.1 µmol), washed twice with PBS, and re-incubated in 10% DMEM for 4 h. Cells were fixed using 4% paraformaldehyde, blocked in PBS-TBH (0.1% Triton X-100, 10% horse serum, and 1% BSA in PBS) for 30 minutes and then incubated in primary and secondary antibodies for 2 h, respectively. Cells were mounted using VECTASHIELD Antifade Mounting Medium with DAPI (H-1200) and imaged using a Leica TCS SP8 confocal microscope.

### Lysotracker Red Stain Assay

Vero cells were seeded at 70–80% confluency on 4-well slides. Cells were transfected with a CIN85-Flag plasmid using Lipofectamine 3000 Transfection kit (Thermo Fisher Scientific) and then 24 h later were infected with WT, ICP0-RF, or ΔICP34.5 (10 PFU/cell). LysoTracker Red stain was added 12 hours post infection for 1 hour at 37 ^0^C (0.1 µmol), washed twice with PBS, and re-incubated in 10% DMEM for 2 h. Cells were fixed using 4% paraformaldehyde, blocked in PBS-TBH (0.1% Triton X-100, 10% horse serum, and 1% BSA in PBS) for 30 minutes and then incubated in primary and secondary antibodies for 2 h, respectively. Cells were mounted using VECTASHIELD Antifade Mounting Medium with DAPI (H-1200) and imaged using a Leica TCS SP8 STED.

### DQ-BSA Assay

Hep-2 cells were seeded on glass slides and infected either 12 h or 4 h with 10 PFU/cell, then stained with 10 µM DQ Red BSA (Thermo Fisher Scientific) for 30 min at 37 ^0^C, then washed with 1% DMEM. Rapamycin was used at 5 uM concentration for a 2 h incubation pretreatment in HEp-2 cells, then washed with 1% DMEM followed by DQ BSA dye staining. The cells were live imaged 4 h post dye staining using Invitrogen™ Attofluor™ Cell Chamber for microscopy (Thermo Fisher Scientific) and a Leica TCS SP8 confocal microscope. Mean Fluorescence Intensity (MFI) was calculated using Image J for 40 cells per sample.

### EV purification and exposure assays

Procedures were as before with minor modifications (57, 58, 110). hTERT-HEL cells were infected at 0.5 PFU/cell for 48 h. Supernatant was subjected to a centrifugation at 5,000 rpm for 5 min to remove cells, and then at 3500 rpm for 20 mins to pellet cell debris. Supernatant was then filtered through a 0.45-μm filter and concentrated using Centricon Plus 70 (100-kDa cutoff) (Millipore). For Western blot analysis EVs were pelleted at 100,000x *g* for 1 h. For EV exposure assays, hTERT-HEL cells were uninfected or infected with WT and ICP0 mutant viruses (0.3 PFU/cell). Supernatant was collected 17 hpi, clarified as earlier and concentrated to 1 mL using Centricon Plus 70 (100-kDa cutoff) (Millipore). Sample was loaded on top of an iodixanol/sucrose gradient ranging from 6 to 18% with a 1.2% increment in the concentration of iodixanol (56–58, 61, 62). Samples were centrifuged in an SW41Ti rotor for 150 min at 200,000 x *g* and 4 ^0^C in a Beckman Coulter OPTIMA XPN-80 ultracentrifuge. Fractions (500 μl) were collected from the top of the gradient, totaling 24 fractions. Fractions were analyzed by Western blot and their purity by plaque assays. Then fractions with EVs (1–8) were concentrated together using a 100-kDa-cutoff filter and washed with phosphate-buffered saline (PBS). EVs were quantified using NTA analysis, then exposed to autologous cells for 12 h in numbers listed in the text. Total RNA was extracted using TRIzol, converted into cDNA and expression of several genes was analyzed by RT-qPCR.

### Nanoparticle tracking analysis (NTA)

Nanoparticle tracking analysis (NTA) was performed using the Malvern Panalytical NanoSight LM10 instrument. NTA software version 3.3 was used to analyze 60-s videos of data collection to obtain the mean, median, and mode of vesicle size and concentration. Results represent the averages of three independent EV isolations.

### mRNA quantification

Our mRNA isolation and quantification protocol is listed elsewhere (58, 111, 112). Briefly, cells were lysed in 1 ml of TRIzol reagent (Life Technologies) and total RNA was extracted using phenol-chlorofom procedures. DNase treatment was performed using Turbo DNase enzyme (Ambion) and cDNA was generated from 1 ug of total RNA per sample using LunaScript® RT Master Mix Kit (Primer-free) (New England Biolabs E3025L). Real-time PCR analyses were performed using SYBR green reagent (Invitrogen) or TaqMan (applied Biosystems) according to the manufacturer’s recommendations. 18S was used for normalization. Predesigned probes (FAM/MGB) for human ISG15, IFIH1, and IL-1β transcripts were obtained through Thermo Fisher Scientific. The primer for ISG56 and 18S are listed elsewhere (57, 110–112).

### Statistical analysis

The p values were calculated using a two-tailed Student’s t-test with a p <0.05 considered significant or a two-way ANOVA with Tukey’s post hoc. Stars represent the following values: * p <0.05, ** p <0.01, *** p < 0.001 and **** p <0.0001. All statistical analyses were performed using at least three biological replicates to ensure reproducibility.

## Acknowledgements

This study was supported by the NIAID R01AI162784, the NIAID R21AI144883, and the NIGMS P20GM130448. Sarah Lasnier is supported by the F31AI186315. The KUMC Leica TCS SP8 STED is supported through the NIH S10 OD 023625.

